# Microbial colonization establishes stratified radial niches that coordinate host epithelial and immune maturation in the colon

**DOI:** 10.64898/2026.07.31.741947

**Authors:** Han Xiao, Jason X Kang, Wai Sinn Soh, Levene W Chua, Hong Sheng Cheng, Damien Chua, William KK Wu, Ho Ko, Yusuf Ali, Nguan Soon Tan, Yi Liu, Joseph Jy Sung, Sunny H Wong

## Abstract

Microbial colonization is essential for intestinal maturation, yet how spatial organization of the microbiome shapes host tissue function remains unresolved. Here, we applied Stereo-seq V2 spatial platform to simultaneously profile the host transcriptome at single-cell resolution and microbial meta-transcriptome at 5 × 5 μm resolution across the proximal, middle, and distal colon of germ-free (GF) mice and mice reconstituted by fecal microbiota transplantation (FMT), integrated with time-course fecal metagenomics. We observed that, four weeks after FMT, microbial colonization established a mature colonic architecture, increased goblet cell number and mucus layer thickness, diversified epithelial lineages, and expanded stem/transit-amplifying, myeloid, and T-cell populations. Metagenomic profiling showed succession from early colonizers to a metabolically mature, short-chain fatty acid (SCFA)-producing community that stabilized by four weeks. Distance-resolved spatial analysis resolved two reproducible strata of colonized microbiota along the radial host-lumen axis, separated at approximately 150 μm. The epithelium-proximal stratum was enriched for mucus-associated taxa such as *Bacteroides thetaiotaomicron*, whereas the luminal stratum harbored fiber-associated taxa such as *Ruminococcus champanellensis*. This radial organization was underpinned by co-occurrence networks of spatial co-localization and co-exclusion. Finally, we identified a butyrate-producing guild that preferentially colonized the epithelium-proximal stratum, localized closer to epithelial and stromal cells, and showed active butyrate-responsive transcriptional activity. Colonization therefore establishes a spatially integrated host–microbiome interface with quantifiable, stratified microbial niches in which location, and not composition alone, coordinates epithelial and immune maturation.

## Introduction

The mammalian intestine harbors one of the most densely populated microbial ecosystems in biology, reaching 10¹¹–10¹² cells per milliliter^1^. This ecosystem is spatially structured along both longitudinal (proximal–distal) and transverse (mucosa-lumen) axes by gradients in oxygen, nutrients, antimicrobial peptides, and signaling pathways that maintain mucus architecture^2^. The mucus layer normally holds the luminal microbiota away from the epithelial surface, and this spatial segregation is central to barrier function and immune homeostasis^3^. Loss of this segregation, permitting bacteria to encroach on the epithelium, is a recurring feature across diverse digestive conditions: the mucus barrier becomes penetrable and bacteria reach the epithelium in inflammatory bowel diseases (IBD)^4^; invasive bacterial biofilms breach the mucus in colorectal cancer (CRC)^5^; and diet-driven perturbations erode the barrier, through fibre deprivation that drives microbial degradation^6^ or dietary emulsifiers that thin the mucus layer and reduce the distance between bacteria and the epithelium^7,8^.

Germ-free (GF) models have established that microbial colonization is essential for intestinal maturation. In the absence of microbiota, GF mice exhibit shortened crypts, reduced goblet cell maturation, a thinner and more penetrable mucus layer, and impaired immune compartment development^3,9^. Colonization restores epithelial proliferation, mucus organization, and immune differentiation, demonstrating that the microbiome gives direct signals to drive structural and immunological development of the gut^3,9,10^. However, these studies rest on bulk or dissociated measurements, and do not resolve locations of the microbes relative to the tissue they act upon.

Recent advances in spatial transcriptomics have revealed that the intestinal epithelium is zonated along crypt–villus and proximal–distal axes^11,12^. The microbiota-induced transcriptional adaptation is spatially restricted rather than globally uniform, with defined intestinal regions acting as hotspots of immune-mediated adaptation^13^. In parallel, imaging approaches such as HiPR-FISH and three-dimensional quantitative microscopy have demonstrated non-random microbial spatial organization and conserved co- localization networks^14,15^. Sequencing-based spatial host–microbiome methods such as SHM-seq and spatial meta-transcriptomics have provided proof-of-principle mapping of host–microbe niches^16,17^. Stereo-seq V2 further enables high-resolution total RNA capture, including non-polyadenylated bacterial transcripts, at near single-cell spatial resolution^18^. What remains missing is a genome-wide, distance-resolved map that places microbial taxa on a continuous axis from the epithelium. Stool-based sequencing does not capture the mucosal ecology, and the relationship between luminal and epithelial ecology following fecal microbiota transplantation (FMT) remains unclear^19^.

Here, we tested the hypothesis that microbial colonization organizes the colon into stratified spatial ecological niches that coordinate epithelial and immune maturation. Using Stereo-seq V2, we profiled host transcriptomes at single-cell resolution and microbial meta-transcriptomes at bin10 (5 × 5 µm) resolution across the proximal, middle, and distal colon of GF mice, and of GF mice reconstituted by FMT from specific pathogen-free (SPF) donors, and sampled 28 days after gavage. This allowed us to ask whether adult intestinal maturation is an emergent property of spatial ecological order rather than of microbial composition or diversity alone.

## Results

### A simultaneous host and microbial map of the colonized colon

To define the spatial landscape of microbial colonization in the colon, we applied random-primer-based Stereo-seq V2 to transverse sections of the proximal (P), middle (M), and distal (D) colon of GF mice and FMT recipients, resolving host transcriptomes at single-cell resolution and microbial meta-transcriptomes at bin10 (5 × 5 μm) resolutions (Fig. 1A).

**Figure 1.**
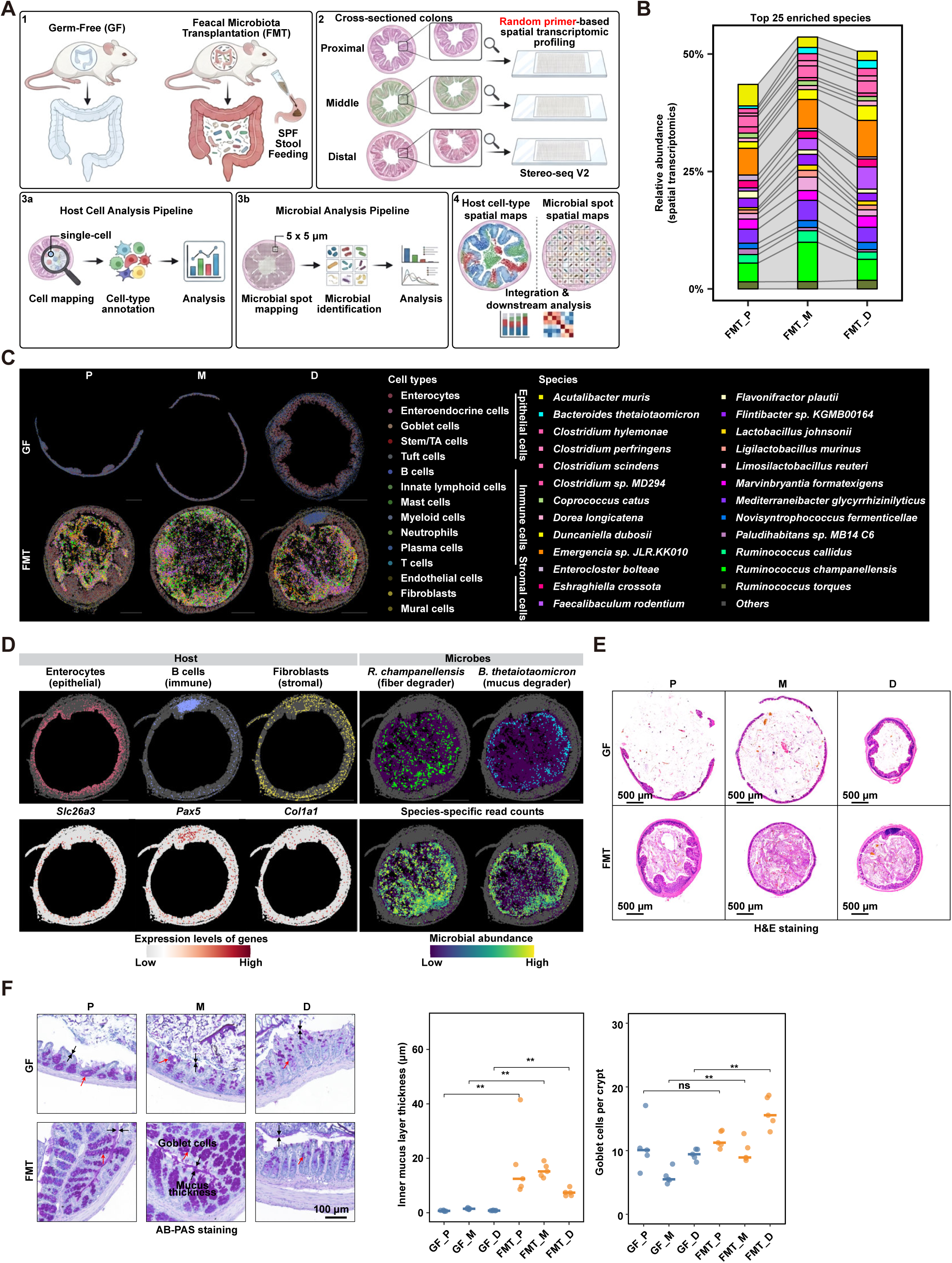
High-resolution spatial transcriptomics reconstructs the host–microbiota landscape of the colon. **(A)** Experimental design for spatial profiling of host and microbial transcriptomes in the proximal (P), middle (M), and distal (D) colon of germ-free (GF) and faecal microbiota– transplanted (FMT) mice using random-primer-based Stereo-seq V2. **(B)** Stacked bar charts showing the relative abundance of the top 25 enriched microbial species across FMT proximal, middle, and distal colon segments. **(C)** Spatial distribution maps of annotated host cell types and microbial species in transverse colon sections of GF and FMT mice. Scale bar: 500 µm. **(D)** Scatter plot showing the spatial expression of representative host marker genes (*Slc26a3* for enterocytes, *Pax5* for B cells, *Col1a1* for fibroblasts), alongside representative microbial signals (*R. champanellensis* as a fiber degrader, *B. thetaiotaomicron* as a mucus degrader). Scale bar: 500 µm. **(E)** Representative H&E-stained transverse colon sections of GF and FMT mice across colonic regions. **(F)** Representative Alcian blue–PAS-stained transverse sections showing regional mucus distribution. Quantification of the inner mucus layer thickness (bottom middle) and goblet cells per crypt (bottom right) across GF and FMT colonic regions. ** *P* < 0.01; ns, not significant; Wilcoxon rank-sum test.

FMT tissues exhibited a high microbial burden, yielding 1,861,283 microbial reads in total (Extended Data Fig. 1A). Following quality control, we identified 100,570 microbial- positive spots. Environment-associated taxa accounted for 0.8% of microbial signals after filtering (Extended Data Fig. 1B; Methods). Because individual microbial spots could contain signals from multiple taxa, spots were further classified as dominant-species or mixed-species spots based on the distribution of species-specific UMI counts (Methods). Overall, 62,253 spots (61.9%) carried a dominant species and were annotated accordingly (Extended Data Fig. 1C, D). The 25 most abundant microbial species (>1% relative abundance) accounted for approximately 49% of signal and maintained highly similar profiles among the three colonic regions (Fig. 1B).

Spatial mapping revealed that microbial signal was overwhelmingly luminal, only rarely crossing the epithelial boundary (Fig. 1C; Extended Data Fig. 1E, F), as expected of an intact barrier. The robustness of cell annotations was further supported by distinct, cell-type-specific transcriptional signatures and their corresponding biological pathway enrichments across colonic segments (Extended Data Fig. 2A-B). Importantly, host and microbial signals both occupied their expected positions: *Slc26a3*, *Pax5*, and *Col1a1* marked enterocytes, B cells, and fibroblasts respectively^20–22^, while the fiber degrader *Ruminococcus champanellensis* preferentially occupied the central lumen and the canonical mucus-foraging *Bacteroides thetaiotaomicron* preferentially localized adjacent to the epithelial surface^23,24^ (Fig. 1D; Extended Data Fig. 1F; Extended Data Fig. 2A).

### Colonization restores colonic architecture and mucus barrier integrity

Colonization promoted structural maturation of the murine intestine. The colons of GF mice exhibited a simplified morphology, with a thinner intestinal wall and shallower, less densely packed crypts compared to FMT mice (Fig. 1E). Alcian blue–PAS staining showed a significantly thicker inner mucus layer and more goblet cells after colonization (Fig. 1F).

### Microbial stabilization follows coordinated ecological succession

We next examined the temporal dynamics of microbial engraftment using shotgun metagenomic sequencing of stool samples collected at Day 3 (D3), Week 1 (1WK), Week 2 (2WK), and Week 4 (4WK) after gavage (Fig. 2A). Alpha diversity was markedly lower at D3, reflecting an early ecological bottleneck, but progressively recovered to approach donor levels by 4WK (Fig. 2B). PCoA and differential abundance analyses showed major compositional remodeling confined to the first two weeks (Fig. 2C, D).

**Figure 2.**
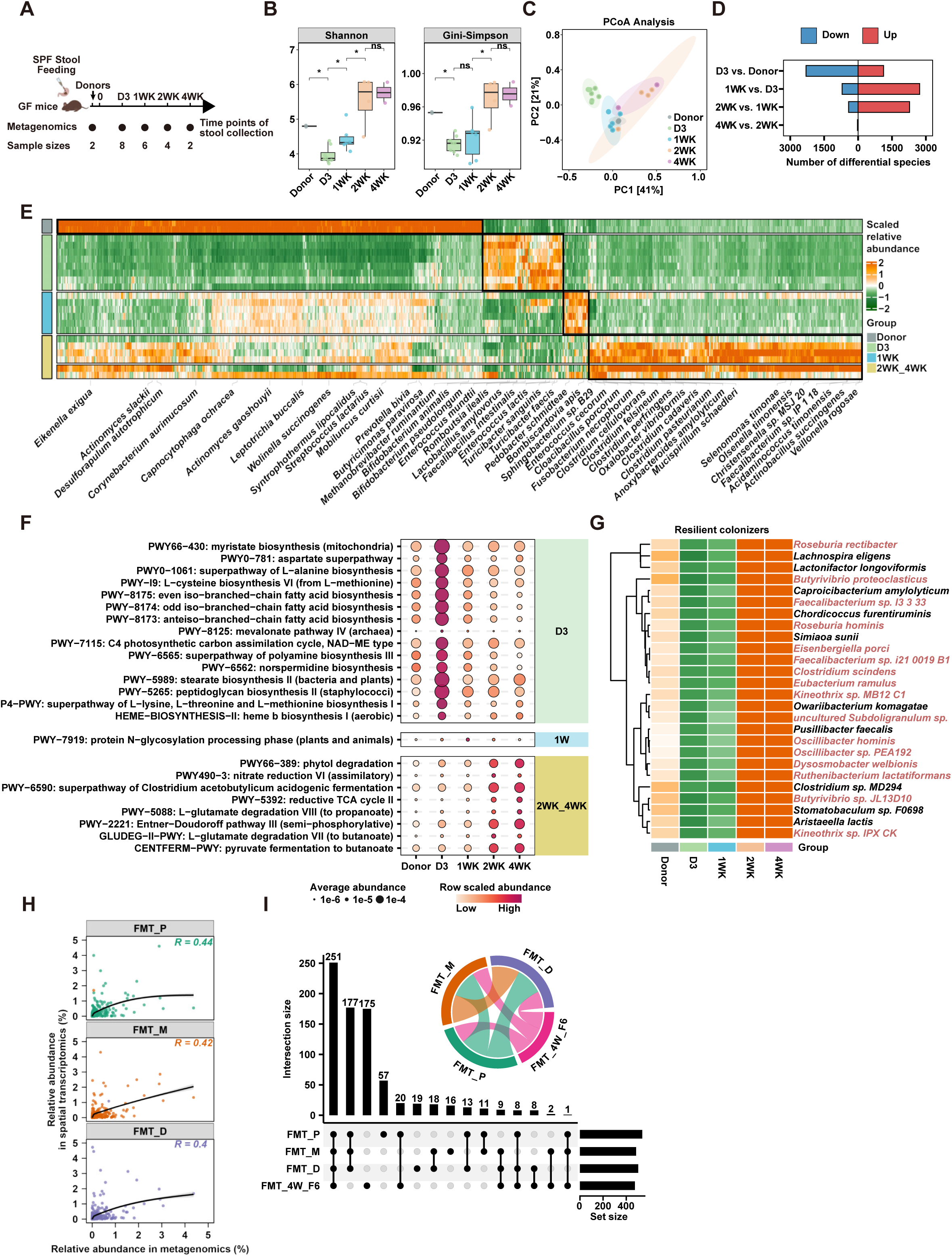
Microbiome maturation follows coordinated taxonomic and functional succession. **(A)** Experimental timeline for tracking microbial colonization in GF mice inoculated with SPF stool. Faecal samples were collected for metagenomics at day 3 (D3), 1 week (1WK), 2 weeks (2WK), and 4 weeks (4WK). **(B)** Boxplots showing the Shannon and Gini-Simpson diversity indices of the gut microbiota in donors and recipient mice across the indicated time points. **(C)** Principal coordinate analysis (PCoA) showing the successional trajectory of the gut microbiota assembly. **(D)** Bar plot detailing the number of differentially abundant microbial species between consecutive time points and compared to the donor. Up, up-regulated genes; Down, down-regulated genes. **(E)** Heatmap showing the longitudinal abundance dynamics of time-point-specific microbial species during the 4-week colonization. **(F)** Dot plot showing the average abundance of differentially time-point-specific metabolic pathways during ecological succession. **(G)** Heatmap showing the abundance dynamics of resilient core colonizers that recovered during the late maturation phase (2WK-4WK). Species with butyrate-producing capacity are highlighted in red. **(H)** Scatter plots showing the Spearman correlation (*R*) between species relative abundance quantified by metagenomics (FMT_4W_F6) and spatial transcriptomics across FMT regions from the same sample. **(I)** Upset plot showing the intersection size of shared microbial species among the proximal, middle, and distal colon, as well as faecal sample.

To characterize the ecological succession, we examined the temporal dynamics of microbial taxa (Extended Data Fig. 3A). Dominant donor taxa including *Ligilactobacillus murinus* and *Adlercreutzia equolifaciens* declined sharply by D3. Glycan-utilizing species, such as *Bacteroides thetaiotaomicron*, *Bacteroides ovatus*, and *Bifidobacterium pseudolongum*, expanded during the first week, followed by the emergence of complex polysaccharide degraders and short-chain fatty acid (SCFA)-producing bacteria, including multiple *Alistipes* and *Clostridium* species, during weeks 2–4 (Extended Data Fig. 3B). Time-point-specific analysis revealed a highly coordinated ecological succession (Fig. 2E).

Further pathway analysis revealed a similar succession pattern (Extended Data Fig. 3C, D). Early colonization (D3) was enriched by amino acid, fatty acid, and peptidoglycan biosynthesis, consistent with rapid microbial proliferation, whereas in weeks 2-4 the microbial community shifted toward fermentative functions, such as pyruvate and L-glutamate fermentation to butanoate, and canonical *Clostridium* acidogenic fermentation (Fig. 2F; Extended Data Fig. 3E). Of the 26 bacterial species that underwent an early decline but recovered upon later maturation (Fig. 2G), 16 (61.5%) species were recognized SCFA producers, principally butyrate producers such as *Roseburia* spp., *Butyrivibrio* spp., and *Faecalibacterium* spp..

### Colonic microbiome is a conserved core community that is regionally fine-tuned

We examined the conservation and regional specialization of the gut microbiota across colonic regions. We observed that fecal and tissue communities were concordant (Spearman R = 0.40–0.44), and species shared between fecal (FMT_4W_F6) and colonic regions (FMT_P/M/D) formed the largest set (Fig. 2H-I).

Between regions, a total of 428 species were shared, representing 79%, 88%, and 85% of taxa (relative abundance >0.01%) detected in proximal, middle, and distal colons respectively (Extended Data Fig. 3F). These included mucus-foraging bacteria such as *Akkermansia muciniphila*, *Bacteroides thetaiotaomicron*, *Ruminococcus torques*, and *Mediterraneibacter gnavus*, whose abundances were differentially tuned across colonic regions (Extended Data Fig. 3G). Despite this overall conservation, region-specific taxa were detected (Fig. 2I; Extended Data Fig. 3F), suggesting that the colon adopts a shared community with fine-tuning to cater for regional adaptation.

### Colonization establishes radial stratification along the host–lumen axis

To resolve microbial organization along the radial host–lumen axis, a spatial dimension that remains largely unexplored, we developed a distance-resolved computational framework integrating cross-sectional spatial meta-transcriptomics with host tissue architecture. For each microbial spot, we computed the minimal distance to the nearest host cell, partitioned the luminal compartment into consecutive, non-overlapping 10-μm bands, and scored every species within each band as the product of its prevalence and normalized expression (Fig. 3A–C; Methods).

**Figure 3.**
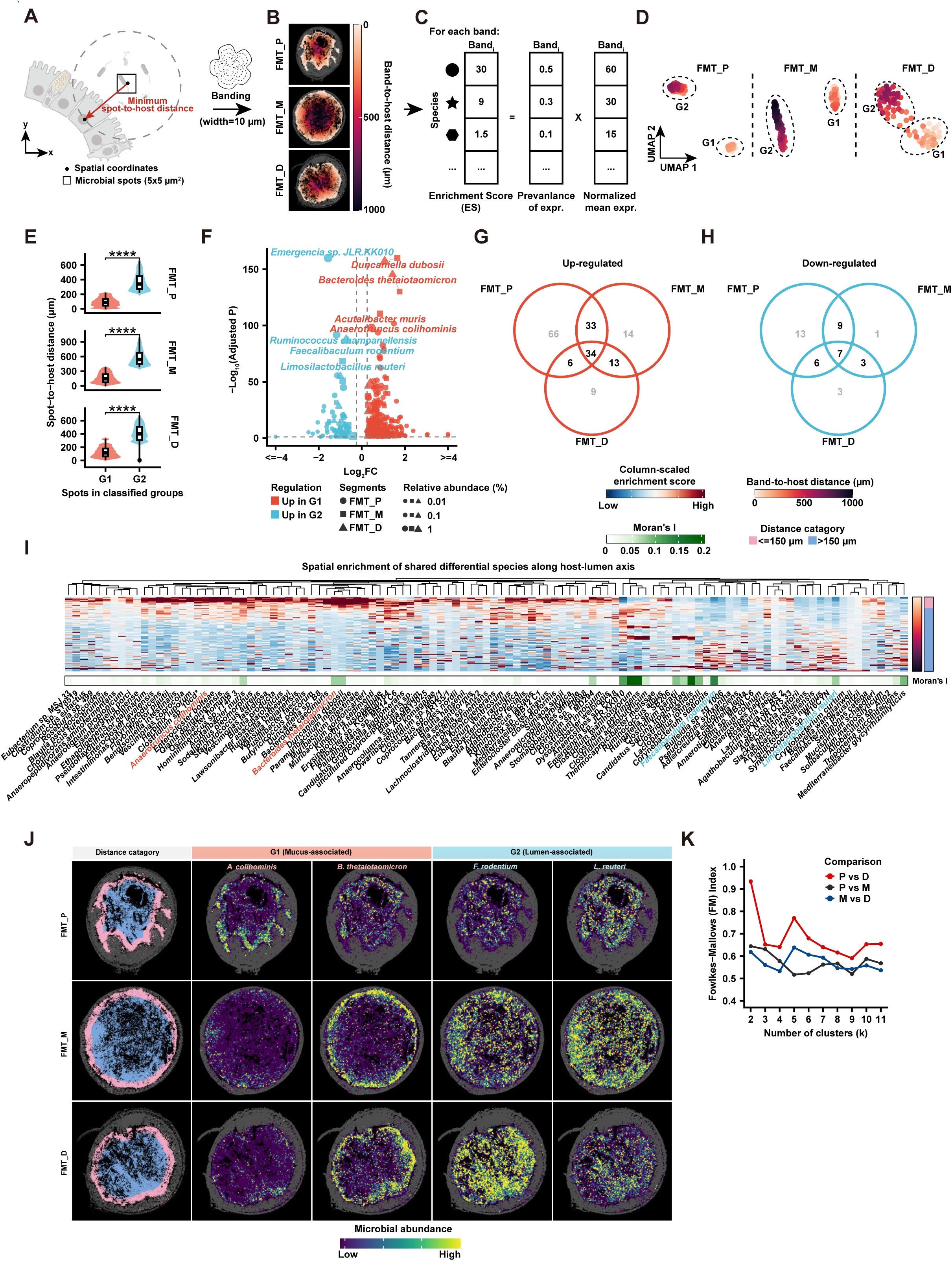
Microbial colonization establishes radial ecological niches along the host–lumen axis. **(A–C)** Computational framework for spatial niche identification. Schematic illustrating the calculation of minimum spot-to-host distance (A), distance-based spatial banding with 10- µm width (B), and the calculation of the species-level enrichment score (ES) for each band (C). **(D)** UMAP embeddings of spatial bands clustered into mucus-associated (G1) and lumen- associated (G2) niches across FMT colon. **(E)** Violin plots comparing spot-to-host distances between G1 and G2 spatial groups across colonic segments. **** *P* < 0.0001; Wilcoxon rank-sum test. **(F)** Volcano plot highlighting the most significantly enriched microbial species between G1 and G2 regions. **(G–H)** Venn diagrams showing the overlap of significantly up-regulated (G1-enriched, mucus-associated) (G) and down-regulated (G2-enriched, lumen-associated) (H). **(I)** Heatmap showing the column-scaled enrichment scores of shared differential species indicated in (G-H) along the host-lumen axis in FMT middle colon. **(J)** Representative spatial maps showing the distinct localization patterns of G1- associated (*A. muris, B. thetaiotaomicron*) and G2-associated (*F. rodentium, L. reuteri*) microbial species in transverse colon sections. Scale bar: 500 µm. **(K)** Dot-line plot of Fowlkes-Mallows (FM) indices comparing the hierarchical clustering of microbial spatial distributions between pairs of colonic segments across different numbers of clusters (k).

UMAP analysis of band-level enrichment profiles identified two distinct spatial clusters, G1 and G2, across every colonic segment (Fig. 3D). G1 lay significantly closer to the epithelium than G2 (mean = 130 μm versus 446 μm, *P* < 0.0001, Fig. 3E), with the transition at approximately 150 μm. This boundary is consistent with the reported thickness of the total murine colonic mucus layer ^25^. Accordingly, G1 and G2 were designated as the mucus-associated and lumen-associated strata.

We observed that the two strata had distinct microbial memberships. Canonical mucus-foraging bacteria, including *Bacteroides thetaiotaomicron*, together with *Duncaniella dubosii*, *Acutalibacter muris*, and *Anaerotruncus colihominis*, were significantly enriched in G1, whereas *Emergencia* sp. JLR.KK010, *Ruminococcus champanellensis*, *Faecalibaculum rodentium*, and *Limosilactobacillus reuteri* preferentially occupied G2 (Fig. 3F; Extended Data Fig. 4A). Across segments, 86 mucus-associated and 25 lumen-associated species showed the same conserved niche preference in two or more segments (Fig. 3G-H). These taxa displayed sharply separated abundance profiles in either side of the 150 μm boundary (Fig. 3I-J).

Apart from radial stratification, microbial localization manifested local clustering. Moran’s *I* identified strong spatial self-aggregation for *Emergencia* sp. JLR.KK010, *Ruminococcus champanellensis*, and *Faecalibaculum rodentium*, as well as previously reported spatial clustering of the *Clostridium* genus^26^, driven by *Clostridium perfringens*, *Clostridium isatidis*, and *Clostridium manihotivorum* (Fig. 3I; Extended Data Fig. 4B–C).

The radial microbial architecture was conserved along the colon. Comparing the hierarchical clustering of the 428 shared species by the Fowlkes–Mallows (FM) index, the distal and proximal colons were more similar, whereas the middle colon showed modestly lower concordance (Fig. 3K), consistent with reports of distinct epithelial programs and host–microbiota interactions in the middle colon^13^. The colonized colon is therefore not a homogeneous luminal community but a radially stratified one, organized by distance from the epithelium and reproducible across its length.

### Spatial co-occurrence networks within the stratified communities

Microbial communities manifest complex ecological networks that shape community structure and function^27,28^. For each microbial spot, we compared the frequency of neighboring species within a 25-μm radius against permuted null distributions, classifying pairs as spatial co-localization or co-exclusion (Extended Data Fig. 5A; Methods).

We identified an average of 1,851 co-localization and 74 co-exclusion species pairs (*P* < 0.01), of which 39 co-localizations and 13 co-exclusions were conserved across all three regions (Fig. 4A, B; Extended Data Fig. 5B, C). Several were ecologically interpretable, including the co-localization between *B. thetaiotaomicron*, a mucin glycan degrader, and *D. dubosii*, a specialized polysaccharide fermenter, suggesting potential metabolic cooperation through carbohydrate cross-feeding. Conversely, *R. champanellensis* emerged as a major co-exclusion hub, spatially segregating from *Acutalibacter muris*, *Clostridium scindens*, *Flavonifractor plautii*, and *Intestinimonas butyriciproducens* (Fig. 4A, B; Extended Data Fig. 5B, C), consistent with a highly specialized ecological niche to spatially segregate from potential competitors.

**Figure 4.**
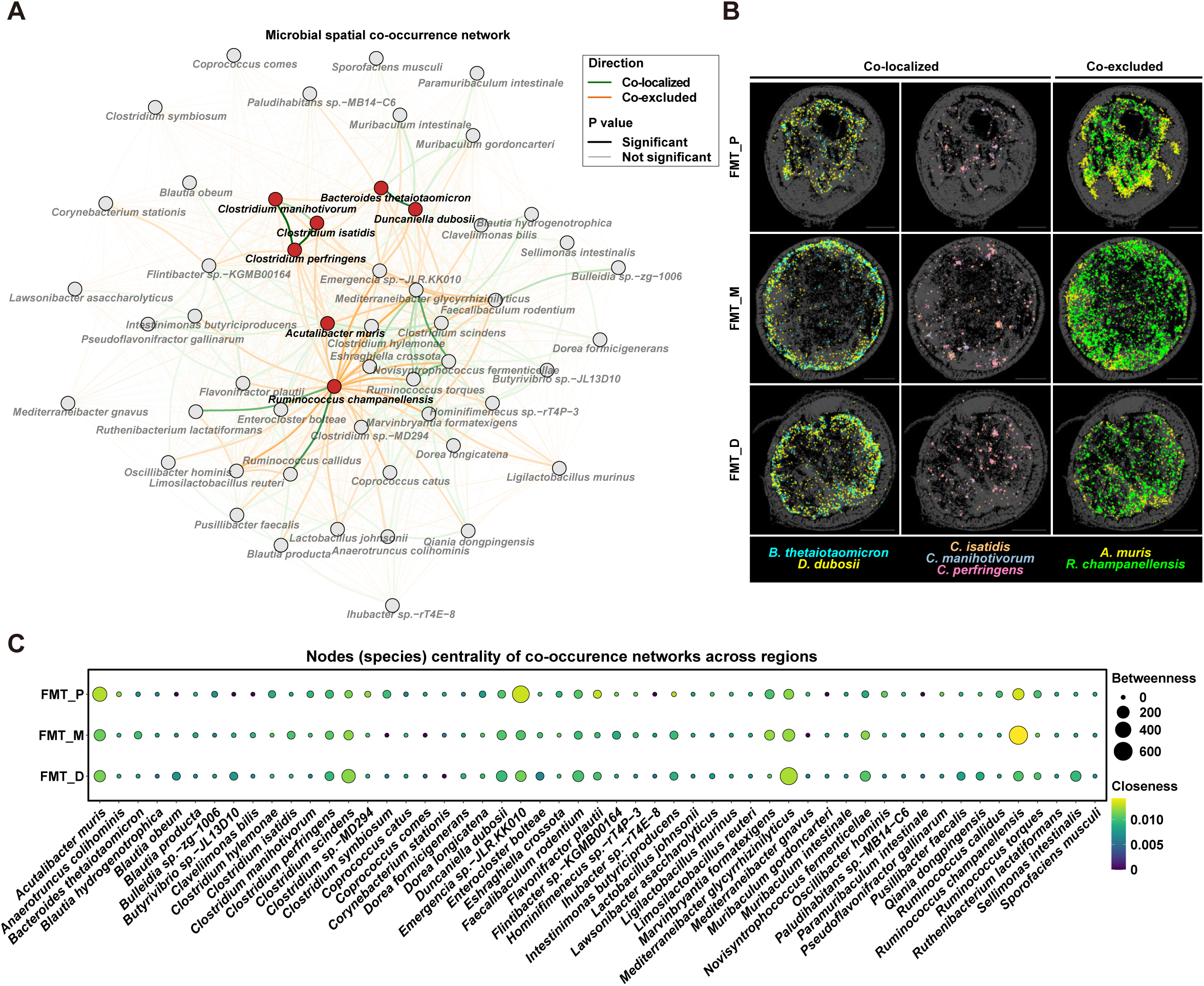
Conserved spatial interaction networks organize microbial communities. **(A)** Spatial co-occurrence network of microbial species in the FMT middle colon. Nodes represent species and edges indicate significant spatial co-localization (green) or co- exclusion (orange). Edge line type denotes statistical significance (*P* < 0.01). **(B)** Spatial mapping of representative co-localized (*B. thetaiotaomicron* and *D. dubosii*; *C. isatidis*, *C. manihotivorum*, and *C. perfringens*) and co-excluded (*A. muris* and *R. champanellensis*) species pairs across colon segments. Scale bar: 500 µm. **(C)** Dot plot showing the betweenness and closeness centrality of microbial species in spatial co-occurrence networks across regions.

The architecture of these networks was itself conserved: betweenness and closeness centrality distributions were closely matched across P, M and D, with no significant regional difference in betweenness (Fig. 4C; Extended Data Fig. 5D). Spatial position in the colonized colon is therefore governed by reproducible interaction rules rather than by regional composition.

### Microbial colonization remodels the host cellular landscape

We next investigated how microbial colonization reshapes host cellular composition and transcriptional programs across the colon. Following quality control, 57,993 high-quality cells were retained for downstream analyses (Extended Data Fig. 6A). Colonization extensively remodelled epithelial, immune, and stromal compartments (Fig. 5A). Compared to GF mice, FMT restored a more balanced distribution of epithelial, immune, and stromal lineages across all colonic regions (Fig. 5A; Extended Data Fig. 6B-C). Epithelial lineage diversity was significantly higher in FMT than GF tissues (*P* < 0.01; Fig. 5B), while immune and stromal compartments exhibited similar trends (Extended Data Fig. 6D-E). At the cell-type level, stem/TA cells, T cells, and myeloid cells expanded most (Fig. 5C, D), and Ki-67 immunostaining confirmed increased proliferation after colonization (*P* = 0.03, Fig. 5D).

**Figure 5.**
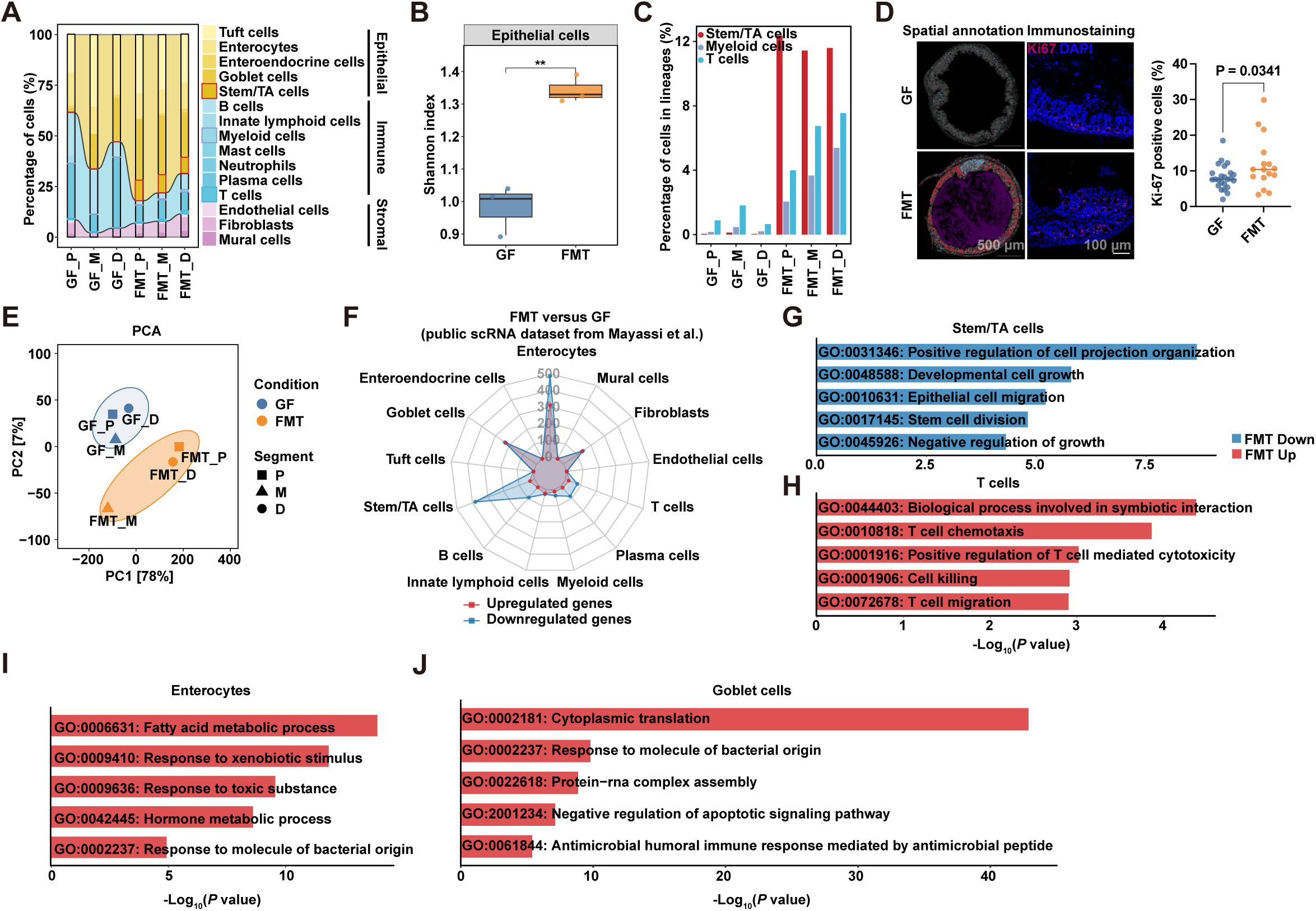
Microbial colonization remodels the cellular and transcriptional landscape of the colon. **(A)** Stacked bar chart showing the relative proportions of distinct cell types. **(B)** Violin plots showing the Shannon diversity index of epithelial cell populations in GF and FMT groups. ** P < 0.01; Student’s t-test. **(C)** Relative percentages of Stem/TA cells, Myeloid cells, and T cells in lineages in GF and FMT groups. **(D)** Spatial distribution of indicated cell types in (C), immunostaining image of Ki67 expression, and dot-line plot showing the percentage of Ki67-positive cells in GF and FMT groups. Student’s t-test. **(E)** Principal component analysis (PCA) of pseudo-bulk gene expressions of spatial transcriptomics across samples. **(F)** Radar plot showing the number of up-regulated and down-regulated genes across distinct cell types using a public scRNA-seq dataset (Mayassi et al.^13^). **(G–J)** Gene Ontology (GO) enrichment of down-regulated genes in Stem/TA cells (G), and up-regulated genes in T cells (H), enterocytes (I), and goblet cells (J).

Principal component analysis (PCA) of pseudo-bulk transcriptomes clearly separated FMT and GF samples, whereas samples from different colonic regions clustered closely together (Fig. 5E; Extended Data Fig. 6F). These findings indicate that microbial colonization is a dominant determinant of host transcriptional program across the colon. Using a condition-matched public single-cell RNA-seq dataset^13^, the largest transcriptional changes occurred in enterocytes, goblet cells, and Stem/TA cells (Fig. 5F). Gene Ontology (GO) enrichment analysis highlighted epithelial renewal in Stem/TA cells, metabolic adaptation in enterocytes, antimicrobial defense in goblet cells, and immune activation in T cells (Fig. 5G–J). Altogether, these findings demonstrate that microbial colonization re-establishes cellular composition, increases lineage diversity, and coordinately reprogram epithelial and immune cell functions toward physiological homeostasis.

### Butyrate-producing guilds occupy the epithelium-proximal stratum

Butyrate is a clear candidate for position-dependent signaling, as it is generated by defined microbial guilds as a key regulator of colonic homeostasis and consumed by colonocytes within the colonic environment^29–31^. Our temporal analyses revealed an enrichment of both butyrate-producing taxa and core butyrate metabolic pathways during late maturation at 2-4 weeks after FMT (Fig. 2E-G; Fig. 6A; Extended Data Fig. 3E). In specific, these canonical butyrate-producing species^32^ exhibited a common trajectory of depletion at D3, rapid expansion between 1-2 weeks, and stabilization by 4 weeks after FMT (Fig. 6B).

**Figure 6.**
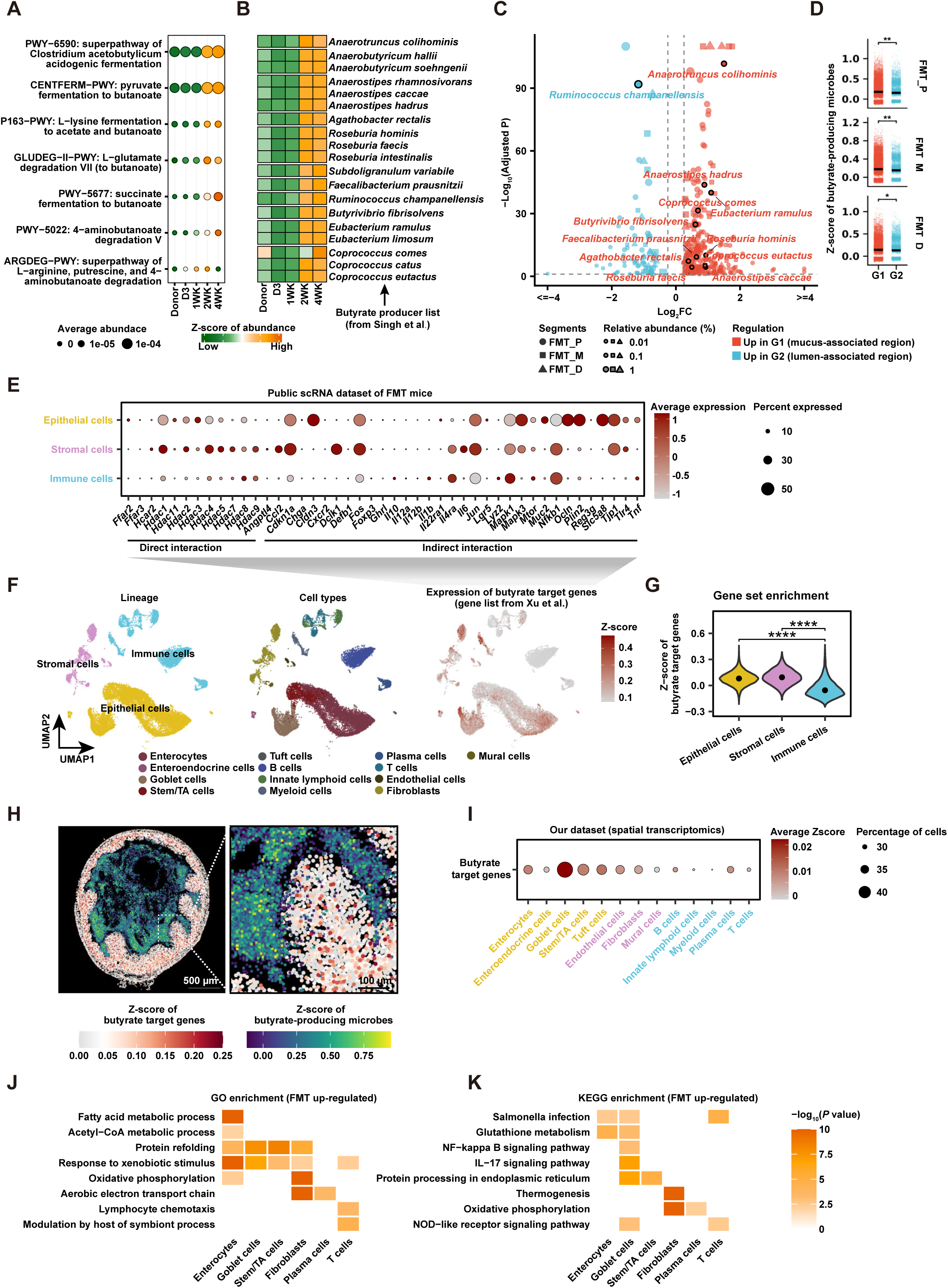
Butyrate-producing microbial guilds establish spatial metabolic crosstalk with the host. **(A)** Dot plot showing the temporal dynamics of core butyrate metabolic pathways from D3 to 4WK post-FMT. Dot size represents the average abundance, and colour indicates the Z-score of abundance (row-scaled). **(B)** Heatmap showing the longitudinal abundance dynamics (column-scaled Z-score) of a curated panel of canonical butyrate-producing microbial species from Singh et al. **(C)** Volcano plot displaying the differential spatial enrichment of butyrate-producing species between the mucus-associated (G1) and lumen-associated (G2) regions across the proximal, middle, and distal colon. **(D)** Violin plots comparing butyrate signature scores of microbial spots between G1 and G2 spatial niches across colonic segments. * *P* < 0.05, *P* < 0.01; Wilcoxon rank-sum test. **(E)** Dot plot showing the expression of representative direct and indirect butyrate target genes across epithelial, stromal, and immune cell lineages, using an independent scRNA- seq dataset (Mayassi et al.^13^). **(F)** UMAP embeddings of host cells from the public scRNA-seq dataset, coloured by lineage (left), annotated cell types (middle), and the Z-score of butyrate target genes (right). **(G)** Violin plot comparing the integrated butyrate target signature Z-score across epithelial, stromal, and immune cell lineages. **** *P* < 0.0001; Wilcoxon rank-sum test. **(H)** Representative spatial maps of the FMT proximal colon section showing the spatial co-localization of host butyrate target gene expression and the abundance of butyrate- producing microbes. **(I)** Dot plot showing the average expression (Z-score) of butyrate target genes across cell types. **(J-K)** Heatmaps showing significantly enriched GO terms (J) and KEGG pathways (K) in indicated cell types.

We investigated whether butyrate producers occupy a specific spatial niche. Differential abundance analysis revealed that most butyrate-producing species were enriched within the mucus-associated region (G1) (Fig. 6C, Extended Data Fig. 7A). Spot-level butyrate signature scores (Z-scores) were significantly higher in G1 than in G2 across all three colonic regions (*P* < 0.01 for FMT_P and FMT_M; *P* < 0.05 for FMT_D), demonstrating that butyrate-producing guilds preferentially colonize the epithelial-proximal mucus niche (Fig. 6D).

Using butyrate target genes curated from the Microbiota–Metabolite–Receptor Database (MMRDB)^33^, we observed that butyrate-responsive genes were predominantly expressed in epithelial and stromal populations (Extended Data Fig. 7B-C). This is consistent with spatial transcriptomic analysis that epithelial lineages displayed the highest butyrate target signature scores (Extended Data Fig. 7D), with representative targets spanning histone deacetylases (*Hdac1* and *Hdac4*), epithelial maturation (*Cdkn1a*), inflammatory responses and repair signaling (*Fos* and *Mapk3*), and barrier maintenance (*Tjp1*) (Fig. 6E). Accordingly, integrated butyrate target signature scores remained significantly enriched within epithelial and stromal cells compared with immune populations (Fig. 6F, G).

To investigate host-microbe spatial coupling, we jointly visualized butyrate-producing bacteria and host butyrate-responsive programs within the same tissue sections. Butyrate producers were concentrated within approximately 100 μm of the epithelial boundary, and the highest butyrate target activities localized to the immediately adjacent enterocytes, goblet cells, and stem/TA cells, together with neighboring stromal populations, including endothelial cells and fibroblasts (Fig. 6H, I), across the three colonic regions (Extended Data Fig. 7E). Gene Ontology (GO) and KEGG enrichment analyses revealed cell type-specific programs downstream of butyrate signaling, including metabolic activity in enterocytes, xenobiotic responses and protein translation in goblet and stem/TA cells, energy metabolism in fibroblasts, and immune activation in T cells (Fig. 6J, K). A functional guild therefore occupies a defined ecological stratum, in this case butyrate-producers, and the host transcriptional response to its product is strongest in the cells that stratum abuts.

## Discussion

Microbial colonization is known to drive intestinal maturation, shaping epithelial renewal, mucus barrier function, and immune development^9,34,35^. Yet, many studies profile microbiomes in bulk and rely on faecal readouts that incompletely reflect mucosal ecology and host-microbe interactions^1,36,37^. Spatial biology studies have highlighted that gut microbes occupy distinct microhabitats (lumen, mucus, crypt-associated niches), but without unifying microbial position with host transcriptional programs in situ^2^.

We address this gap by combining high-resolution spatial host–microbial transcriptomics with orthogonal microbiome profiling. Stereo-seq V2, using random-priming total RNA capture, enables simultaneous detection of host and microbial transcripts at cellular resolution, overcoming the poly(A)-dependence that limits other spatial methods^18^. In contrast to imaging-based taxonomic mapping^38–40^, this yields a sequencing-based spatial meta-transcriptome that can be analysed with distance-resolved statistics, co-occurrence inference, and direct alignment to host states within the same sections.

A central conceptual advance of this study is that colonization establishes distance-defined ecological niches along the host–lumen axis. Classical models emphasize a bilayered mucus barrier in which a dense inner mucus layer excludes bacteria under homeostatic conditions^41,42^, with microbial penetration occurring during inflammatory states such as colitis^4^. Our data show reproducible, species-specific positioning across radial distances from the epithelium, with significant spatial clustering among taxa. This supports a model in which the colon behaves as a graded biophysical and metabolic landscape, where communities distribute along physicochemical gradients rather than a simple binary mucus boundary. The organization of functional guilds is biologically plausible given known host-microbe interactions: colonocyte metabolism and crypt architecture buffer stem and progenitor cells from butyrate exposure^43^, and dietary depletion of microbiota-accessible carbohydrates erodes the mucus barrier and increases microbial proximity to the epithelium^6,39^.

Within this framework, our results reinforce that colonization promotes mucosal maturation, with restored epithelial architecture, strengthened mucus barrier features, and transcriptional programs consistent with epithelial homeostasis and immune maturation^9,11^. Along the longitudinal axis, we find broad continuity of shared taxa with regionally tuned abundance, and the divergent spatial organization resonates with recent mapping identifying microbiota-driven, spatially restricted adaptation in the mid-colon^13^. Co-occurrence networks suggest that microbial communities are not only stratified, but are also organized into interaction architectures^38,40^.

Several strengths increase the value of this work: (i) unified measurement of host and microbes in the same tissue section at high spatial resolution, (ii) quantitative distance-resolved mapping of microbial localization, and (iii) integration with host programs at the host-microbe interface at a cellular level. This positions intestinal maturation as a property of spatial ecological order rather than composition or diversity alone. Limitations include the focus on a single late time-point (Day 28), so the order in which stratification and tissue maturation emerge cannot be determined. Meta-transcriptomic signal reflects activity-weighted RNA abundance rather than absolute biomass, and the use of complex stool-derived communities preclude causal attribution to individual taxa. Furthermore, the butyrate coupling was inferred from producer taxa and target-gene programs rather than directly measured, so producer-responder relationships are positional rather than demonstrated. Taxonomic assignment from short reads is imperfect at species level and the congener-level claims warrant orthogonal confirmation. These limitations should motivate future studies incorporating temporal sampling, orthogonal imaging validation, spatial metabolite mapping, and defined consortia to test causality.

The framework developed here extends directly to contexts where spatial barrier failure drives pathology. In ulcerative colitis, bacteria can penetrate the inner mucus layer to encroach on the epithelium^4^, while in colorectal cancer, invasive polymicrobial biofilms and ecological disruptions have been identified^5^. Similarly, fiber depletion can erode mucus and shift microbial ecology^39^. In each of these, the measurable variable our framework supplies, distance from the epithelium, is precisely the variable that composition-based profiling discards.

In conclusion, we provide a spatially resolved model of colonization-driven maturation in which engraftment produces structured radial and regional niches, conserved interaction architectures, and coordinated epithelial and immune programs. This offers a quantitative blueprint for asking not only which microbes are present in the gut, but where they act upon the host.

## Methods

### Experimental model and study participant details

#### Mice

Six-week-old germ-free (GF) C57BL/6 mice of both sexes were maintained in sterile flexible-film isolators or germ-free individually ventilated cages at the LKCMedicine NTU animal research facility. Mice were housed under a 12-h light/dark cycle with ad libitum access to autoclaved chow and sterile water. Faecal microbiota-transplanted (FMT) mice were generated from age-matched GF mice by oral gavage with a complex stool-derived microbial community from specific pathogen-free (SPF) C57BL/6 donor mice. Unless otherwise stated, mice were sampled 28 days after colonization. Proximal (P), middle (M), and distal (D) colon segments were collected from GF and FMT animals for matched histological, spatial transcriptomic, and microbial analyses. Both male and female mice were included; sex-specific effects were not the primary endpoint, and the study was not powered for sex-stratified analysis.

All animal experiments were approved by the Institutional Animal Care and Use Committee (IACUC) of Nanyang Technological University (approval no. A22065) and were performed in accordance with institutional animal care guidelines.

#### Fecal microbiota transplantation and stool sampling

Fresh donor stool was collected from SPF C57BL/6 mice, suspended in sterile reduced phosphate-buffered saline, homogenized, and clarified to remove large particulates. Recipient GF mice received the stool suspension by oral gavage at the start of the experiment. Colonized mice were monitored after gavage and maintained using procedures designed to prevent cross-contamination between GF and colonized groups. Stool pellets were collected longitudinally at multiple post-colonization time points, including Day 3 (D3), Week 1 (1WK), Week 2 (2WK), and Week 4 (4WK), snap-frozen, and stored at -80°C until DNA extraction. Shotgun metagenomic sequencing was performed on stool samples to evaluate engraftment dynamics, ecological stabilization, taxonomic composition, and functional guild representation.

#### Colon tissue collection and regional sampling

At the experimental endpoint, mice were euthanized according to approved institutional procedures. The colon was excised and divided into proximal, middle, and distal segments. Transverse sections from each segment were processed for Stereo-seq V2 spatial host-microbial transcriptomics, and adjacent or matched sections were processed for histology. Tissue orientation was preserved during embedding and sectioning to retain the host-lumen axis and support distance-resolved analyses from the epithelial boundary into the luminal cavity.

## Method details

### Histology and mucus layer quantification

Colon tissues were fixed in Carnoy’s solution, embedded, sectioned, and stained with Alcian blue-periodic acid Schiff (AB-PAS) to visualize acidic and neutral mucins and assess goblet-cell-associated mucus architecture. Brightfield images were acquired using consistent acquisition settings within each staining batch. Mucus thickness was quantified by measuring the distance from the epithelial surface to the luminal edge of the stained mucus layer at multiple evenly spaced positions per section. Measurements were performed in ImageJ by investigators blinded to the experimental group, where possible. Section-level means were used to compare the GF and FMT groups.

### DNA extraction and shotgun sequencing

DNA was extracted from donor and recipient stool samples using a bead-beating-based microbial DNA extraction procedure optimized for bacterial community profiling. Shotgun metagenomic libraries were prepared, sequenced, and quality-filtered to remove adapters.

### Spatial transcriptomics and meta-transcriptomics sequencing

Spatial transcriptomic profiling of host and spatial meta-transcriptomic profiling of microbes was simultaneously performed using the random-primer-based Stereo-seq V2 OMNI FFPE platform. Transverse colon sections were mounted on Stereo-seq OMNI FFPE capture chips containing spatially barcoded DNA nanoballs and processed for tissue permeabilization, reverse transcription, cDNA amplification, library construction, and sequencing according to the manufacturer’s protocol with modifications for total RNA capture.

### Metagenomics data processing

Raw paired-end reads were quality-filtered using KneadData (v0.12.0) with Trimmomatic (v0.39)^44^. Host and contaminant reads were removed by alignment against the human (hg39), mouse (C57BL/6NJ), and SILVA rRNA databases^45^ using Bowtie2 (v2.4.1, ’--very-sensitive --dovetail’)^46^. High-quality non-host reads were taxonomically profiled using Kraken2 (v2.1.5) with a custom standard database (k2_pluspf_20250402), and species-level relative abundances were estimated using Bracken (v3.0.1)^47^. Functional profiling was performed using HUMAnN (v4.0.0.alpha.1)^48^ with MetaPhlAn (v4.2.2)^49^, and MetaCyc pathway abundances were converted to relative abundances. Species and pathways with a prevalence < 10% across all samples were excluded from downstream analyses. Taxonomic and functional diversity analyses were performed in R (v4.4.3) using phyloseq (v1.50.0)^50^ and vegan (v2.7.1). Alpha diversity was calculated after rarefaction to the minimum sequencing depth and compared using two-sided Wilcoxon rank-sum tests. Beta diversity was assessed by principal coordinate analysis (PCoA) based on Bray-Curtis dissimilarity.

Differentially abundant species were identified using DESeq2 (v1.46.0)^51^ (absolute fold change > 1.5, P < 0.05, Benjamini–Hochberg adjusted P < 0.1), whereas differential pathways were identified using MaAsLin2 (v1.20.0)^52^ with linear models applied to log-transformed relative abundances (absolute coefficient > log_2_(1.5), P < 0.05, a BH-adjusted P < 0.1). Because the 2WK and 4WK groups exhibited highly similar community compositions, they were combined as a single late-stage (2WK_4WK) group. Time-point-specific species and pathways were defined as features significantly enriched in one group relative to each of the other three groups across all pairwise comparisons. Resilient microbial colonizers were defined as species enriched in both the Donor and 2WK_4WK groups relative to D3 and 1WK, showing no significant difference between Donor and 2WK_4WK, and a mean relative abundance > 0.01% in both groups. Heatmaps and Upset plots were generated using ComplexHeatmap (v2.22.0)^53^, UpSetR (v1.4.0)^54^, and eulerr (v7.0.4).

### Spatial transcriptomics data processing

Raw sequencing reads and the matched ssDNA image generated from the Stereo-seq V2 assays were processed with SAW (v8.1.3, count)^55^ following the Stereo-seq OMNI for FFPE workflow^18^. Libraries (Stereo-seq N FFPE V1.0 kit) were sequenced paired-end (PE75_25+59): read 1 (25 nt) carries the coordinate identity (CID) spatial barcode and read 2 (59 nt) carries a 6-nt molecular identifier (MID/UMI) followed by 53 nt of cDNA. CIDs were mapped to the chip mask (barcodeToPos.h5, ≤1 mismatch) to assign each read to its spatial coordinate. After adapter and poly-A trimming, cDNA reads were aligned to the mouse genome GRCm38 (Ensembl, GCA_000001635.8, with matched annotation) using bcSTAR (v3.2.4, based on STAR 2.7.2b)^56^. Uniquely mapped reads and the best-scoring alignment of multi-mapped reads were retained and counted. Reads sharing the same CID, MID and gene were collapsed to remove PCR duplicates. The ssDNA image was registered to the chip and nuclei were segmented; nuclear boundaries were then expanded by ∼10 pixels using a Euclidean distance transform to approximate whole-cell (membrane) outlines. Deduplicated molecules (UMIs) were assigned to cells or spatial bins by their coordinates and summed per gene to generate GEF/GEM expression matrices at single-cell (cellbin) resolution.

The resulting outputs were further imported into StereoMap (v4.1.2) for manual demarcation of tissue boundaries within each segment, generating tissue-region coordinate-mapped spatial gene expression matrices. To ensure high-fidelity of cell transcriptome profiling, low-quality cells were discarded from downstream analysis; only cells with 50 to 500 detected genes and a mitochondrial read fraction of < 10% were retained.

### Spatial meta-transcriptomics data processing

Microbial signal was profiled in the same run using SAW’s microorganism-detection module (--microorganism-detect), which operates on host-unmapped reads^18^. Reads not aligned to the mouse genome by STAR underwent a second host-removal step with Bowtie2 (v2.2.5; --very-sensitive-local --no-unal, single-end)^46^ against the same GRCm38 index; only reads unmapped in both steps were retained as candidate microbial reads. These were taxonomically classified with Kraken2 (v2.1.5)^57^against the prebuilt PlusPF database (k2_pluspf_20250714) using exact k-mer matching and lowest-common-ancestor assignment. Because each classified read retains its spatial coordinate and MID, taxonomic calls were projected back onto the chip and aggregated at each taxonomic rank into spatially resolved microbial expression matrices.

Microbial spots were defined at a Bin10 resolution (5 × 5 μm). Initial quality control excluded microbial spots with total UMI counts < 10. The top 10 species in GF samples accounted for 63% of all microbial reads (2,727) and were predominantly composed of non-gut-associated taxa, (Extended Data Fig. 1A). They were further defined as environment-associated taxa, and their proportional abundances were subsequently calculated within FMT samples and used to filter microbial spots for quality control. High-quality microbial spots were retained for downstream analysis based on the following thresholds: total UMI counts ≥ 10, detected species ranging from 1 to 30, and an environmental contaminant proportion ≤ 10%.

### Cell type annotation

An annotated murine colon scRNA-seq reference dataset^13^ was utilized to deconvolute the spatial transcriptomics data. For GF samples, segment-specific concordance (proximal, middle, and distal) was strictly maintained between the spatial queries and the scRNA-seq references. However, as region-specific reference data for FMT conditions were limited, all FMT spatial segments were deconvoluted using the FMT middle colon scRNA-seq reference. Spatial mapping was performed using cell2location (v0.1.4)^58^. Reference cell type signatures were estimated by training a negative binomial regression model on the scRNA-seq counts (max_epochs=1500, batch_size=2000, train_size=1, lr=0.002). The spatial mapping model was subsequently trained on the spatial data (max_epochs=2000, train_size=1, lr=0.002, detection_alpha = 300, N_cells_per_location = 2) to estimate absolute cell type abundances across multiple hierarchical annotation levels. Deconvolution weights were integrated into spatial objects using the R package Seurat (v5.4.0)^59^. Cells were discretely annotated by assigning the cell type with the maximum predicted probability weight.

### Cell type proportions and diversity analysis

Cell type proportions were quantified globally and separately within distinct lineages (epithelial, immune, and stromal) based on annotated results. The Shannon diversity index of cell types was subsequently calculated within each lineage for each sample. Diversity differences between the FMT and GF groups within anatomically matched segments were evaluated using the Student’s t-test. Alluvial diagrams were visualized using the R package ggalluvial (v0.12.6)^60^.

### Pseudo-bulk analysis of spatial and single-cell transcriptomics

Spatial and single-cell transcriptomics data were aggregated into pseudo-bulk expression profiles using the PseudobulkExpression function (layer = ’counts’, method = ’aggregate’, group.by = ’Lineage_HiRes’) in the R package Seurat (v5.4.0)^59^. Raw counts were aggregated within each cell type and normalized to counts per million (CPM). For individual samples, the generated CPM matrices were further averaged across all cell types to generate the sample-level gene pseudo-bulk expression matrices. Concordance between the spatial and scRNA transcriptomic datasets was determined using Spearman rank correlation coefficients. Principal component analysis (PCA) was performed on the pseudo-bulk matrices following the removal of zero-variance genes. Heatmaps were visualized using the R package pheatmap (v1.0.13).

### Differential expression analysis of spatial and single-cell transcriptomics

Differential gene expression analysis between the FMT and GF groups within anatomically matched segments was conducted using the FindMarkers function (test.use = ’wilcox’, min.pct = 0.01) in the Seurat package (v5.4.0)^59^. For spatial transcriptomics, significantly up-regulated and down-regulated genes were defined using an absolute threshold of log2 fold-change > 0.1 and P < 0.01. For scRNA transcriptomics, significantly up-regulated and down-regulated genes were defined by an absolute log_2_ fold-change > 0.25 and a Bonferroni-adjusted P < 0.05. Radar charts showing shared up-regulated genes across spatial segments were visualized using the R package fmsb (v0.7.6).

### Functional enrichment analysis

Up-regulated DEGs were subjected to Gene Ontology (GO) biological process and Kyoto Encyclopedia of Genes and Genomes (KEGG) pathway enrichment analyses using the R package clusterProfiler (v4.14.6)^61^ and the mouse genome annotation database (org.Mm.eg.db). Significantly enriched terms were defined by a P < 0.05 and a Benjamini-Hochberg adjusted P < 0.1. Enrichment heatmaps were visualized using the R package pheatmap (v1.0.13).

### Taxonomic annotation of microbial spots

Taxonomic annotation of microbial spots was performed using a combinatorial approach based on the standard deviation (SD) and absolute read counts of detected species within each microbial spot. The SD of species counts was calculated for each microbial spot. Based on a bimodal SD distribution, a threshold of 0.6 was utilized to partition spots into dominant-species and mixed-species categories. Dominant-species spots were defined by an SD > 0.6, absolute counts of the most enriched species ≥ 3, and a count difference of ≥ 1 between the most enriched and second-most enriched species. Spots failing to meet these combinatorial criteria were classified as mixed-species spots. For each dominant-species spot, taxonomy was assigned as the species of the most enriched one, while mixed-species spots were uniformly annotated as "Others".

### Microbial spot-to-host distance analysis

Spatial coordinates for host cells and microbial spots were extracted. Euclidean distances between all microbial spots and host cells were calculated and subsequently converted to physical distances (µm). Given the negligible microbial penetration into the tissue region within our dataset, the spot-to-host distance for each microbial spot was defined as the minimum distance to the nearest host cell.

### Identification of mucus-associated and lumen-associated regions

Microbial spots were classified based on their spot-to-host distances, as defined above. The continuous distance gradient was partitioned into non-overlapping 10-µm spatial bands across the proximal, middle, and distal colon segments. A species-level enrichment score (ES) was calculated for each spatial band as the product of expression proportion and normalized mean expression. The resulting ES matrix was log_2_- transformed with a pseudo-count of 1e-6. Principal component analysis (PCA) was performed on the transformed matrix, and the top 10 principal components were utilized to construct a neighbourhood graph. Uniform Manifold Approximation and Projection (UMAP) was subsequently applied for dimensionality reduction using the R package Seurat (v5.4.0)^59^. Based on the UMAP embedding, spatial bands were clustered into two distinct groups (G1 and G2). Spot-to-host distances between the groups were evaluated using a two-sided Wilcoxon rank-sum test. Consistent with the established 110–150 µm thickness of the murine colonic mucus layer^25^, Group G1 (mean distance = 130 µm) was designated as the mucus-associated region, whereas Group G2 (mean distance = 446 µm) was defined as the lumen-associated region.

### Differential species abundance analysis between mucus-associated and lumen-associated regions

Differential species abundance analysis between the mucus-associated (G1) and lumen-associated (G2) regions was performed independently within each segment. The analysis was performed using the FindMarkers function in the R package Seurat (v5.4.0)^59^, retaining complete feature statistics. Mucus-associated and lumen-associated species were defined as taxa significantly enriched in their respective spatial microenvironments. Statistical significance was defined by an absolute log_2_ fold-change > 0.25, P < 0.05, and a Bonferroni-adjusted P < 0.1. Volcano plot was used to visualise the adjusted P values, fold changes, and relative abundances of species, and specific core target species were annotated using the R package ggrepel (v0.9.8).

### Spatial autocorrelation analysis of microbial features

Spatial autocorrelation of species features was assessed for each sample using Seurat (v5.4.0). The top 2,000 highly variable features following SCTransform were first identified with the variance-stabilizing transformation (VST) method implemented in FindVariableFeatures. Spatial autocorrelation was then quantified by calculating Moran’s *I* for these features using FindSpatiallyVariableFeatures with selection.method = "moransi". Species features with higher Moran’s *I* values were considered to exhibit stronger spatial clustering.

### Comparison analysis of microbial spatial distributions across distinct segments

Enrichment scores of 428 shared species across the proximal, middle, and distal colon segments were extracted across all spatial bands. For individual taxa, enrichment scores were centred and scaled using Z-score normalization across all the bands. Based on the normalized distribution profiles, hierarchical clustering was performed to group species with concordant spatial patterns, and independent dendrograms were constructed for each segment. Pairwise topological comparisons of the resulting dendrograms were conducted. Dendrogram similarity was quantified across a range of cluster numbers (k = 2 to 11) using the Fowlkes-Mallows (FM) index, which evaluates hierarchical structural similarity by computing the geometric mean of the probability that a pair of species co-clustered in one dendrogram is concurrently co-clustered in the comparator dendrogram. Clustering evaluations and topological comparisons were performed using the R package dendextend (v1.19.1)^62^.

### Identification of microbial spatial co-occurrence events

Spatial coordinates of taxonomically annotated microbial spots were extracted. Local spatial co-occurrence was quantified using a distance-weighted neighborhood scoring algorithm across a predefined spatial radius (*r*). For each microbial spot, spatial neighbors within the specified radius were identified using the get.knnx function in the R package FNN (v1.1.4.1). Co-occurrence frequencies between taxa were weighted using a Gaussian decay function formalized as

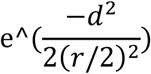

where *d* denotes the Euclidean physical distance between spots and (*r*) denotes the 25- µm radius. An empirical null distribution was generated by randomly permuting the spatial taxonomic labels 100 times. Symmetric Z-scores and two-tailed P-values were calculated by comparing the absolute deviations of observed co-occurrence scores against the mean of the permuted null distribution. Significant spatial co-occurrence events were defined by P < 0.01, with species pairs classified as spatial co-localization (Z-score > 0) or co-exclusion (Z-score < 0).

### Topology analysis of microbial spatial co-occurrence network

To investigate the robustness of co-occurrence events across segments, topological analysis of the constructed microbial spatial co-occurrence networks was performed. Identified significant spatial co-occurrence events were utilized to construct undirected, weighted networks. Network construction and topological calculations were performed using the R package igraph (v.1.5.1). The Louvain clustering algorithm, weighted by absolute Z-scores, was applied to partition the networks into distinct co-occurrence modules. Network centrality metrics, including betweenness centrality and closeness centrality, were computed for individual microbial nodes. The co-occurrence networks were subsequently visualized using the Fruchterman-Reingold layout algorithm weighted by log-transformed absolute Z-scores, with edges coloured by correlation directionality.

### Scoring of expression of butyrate-producing microbes and butyrate target genes

A list of butyrate target genes, including direct and indirect interactions, was retrieved from a custom Microbiota-Metabolite-Receptor Database (MMRDB) from a published study^33^. Signature scores of butyrate target genes in individual cells were calculated using the AddModuleScore function in the R package Seurat (v5.4.0)^59^. A list of classic butyrate-producing species was retrieved from a published review^32^. Similarly, signature scores of butyrate-producing microbes in each microbial spot were computed using the AddModuleScore function. All host or microbial datasets were initially merged to construct a unified background set prior to scoring. Differences in signature scores between groups were evaluated using the Wilcoxon rank-sum test. Spatial distributions of host and microbial signature scores were visualized using custom R functions.

### Quantification and statistical analysis

Statistical analyses were conducted using R (v4.4.3) and Python (v3.9.23). The type of test method used for statistical analysis is specified in the text where the results are described, and details for the test are explained in the relevant figure legend and methods section. Statistical significance was considered when P value < 0.05. ∗, ∗∗, ∗∗∗ and ∗∗∗∗ denote P < 0.05, P < 0.01, P < 0.001 and P < 0.0001, respectively.

## Acknowledgements

The authors would like to acknowledge Junhou HUI, Hui CHEN, Min JIAN and Jingxiang Zhang for their assistance with the data analysis.

## Author Contributions

SHW and JXK conceived and designed the study. HX led the bioinformatics analysis and interpretation of the data. JXK, WSS, LWC, and YL contributed to experimental design, execution, and analysis. HSC, DC, WKKW, HK, YA, and NST provided scientific and intellectual input. SHW, JJYS, LY, and NST provided essential resources and study materials. HX and SHW drafted the manuscript. All authors critically reviewed and approved the final manuscript and agreed to be accountable for the integrity of the work.

## Data availability

The datasets generated and analyzed during the current study are available from the corresponding author upon reasonable request.

---

**Extended Data Fig. 1.**
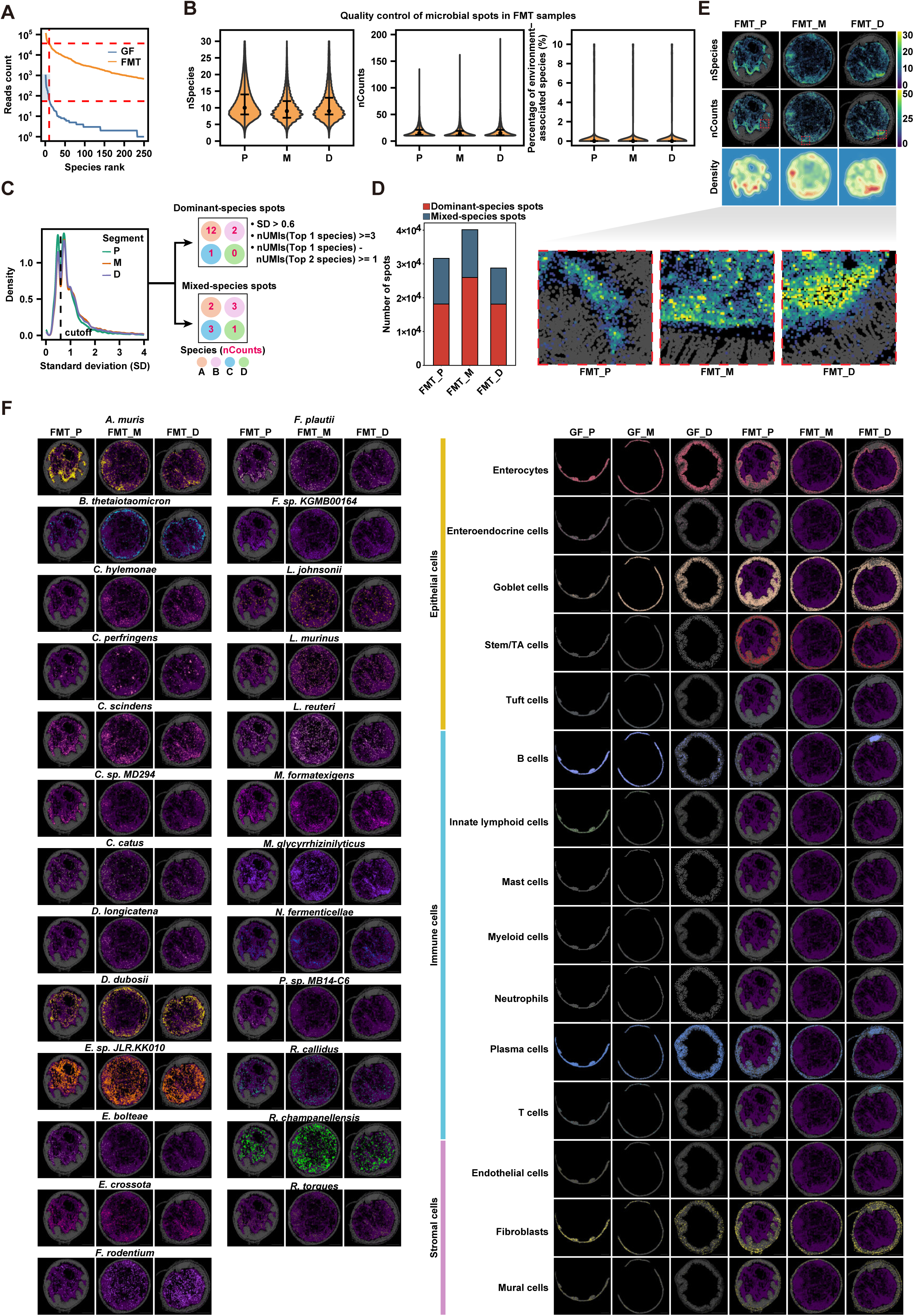
Quality control and microbial spot annotation of spatial microbiome. **(A)** Curves showing ranked total microbial read counts in GF and FMT samples. The top 10 species detected in GF samples are indicated by the red dashed lines and defined as environment-associated species. **(B)** Violin plots showing quality control metrics of microbial spots in FMT samples across segments, including the number of detected species (left), total UMI counts (middle), and percentage of environment-associated species (right). **(C)** Density plot showing the distribution of the standard deviation (SD) of UMI counts across species within microbial spots for each colonic region. A cutoff of SD = 0.6 was used to classify spots. A schematic illustrates the criteria for classifying dominant-species spots versus mixed-species spots based on SD and UMI counts. **(D)** Bar chart showing the number of classified dominant-species and mixed-species spots across FMT colonic segments. **(E)** Spatial maps showing the number of detected species (top) and total UMI counts (middle) and density (bottom) of microbial spots across FMT colonic regions. Scale bar: 500 µm. **(F)** Spatial distribution maps of the top 25 enriched microbial species indicated in Fig. 1B (left) and annotated cell types (right). Scale bar: 500 µm.

**Extended Data Fig. 2.**
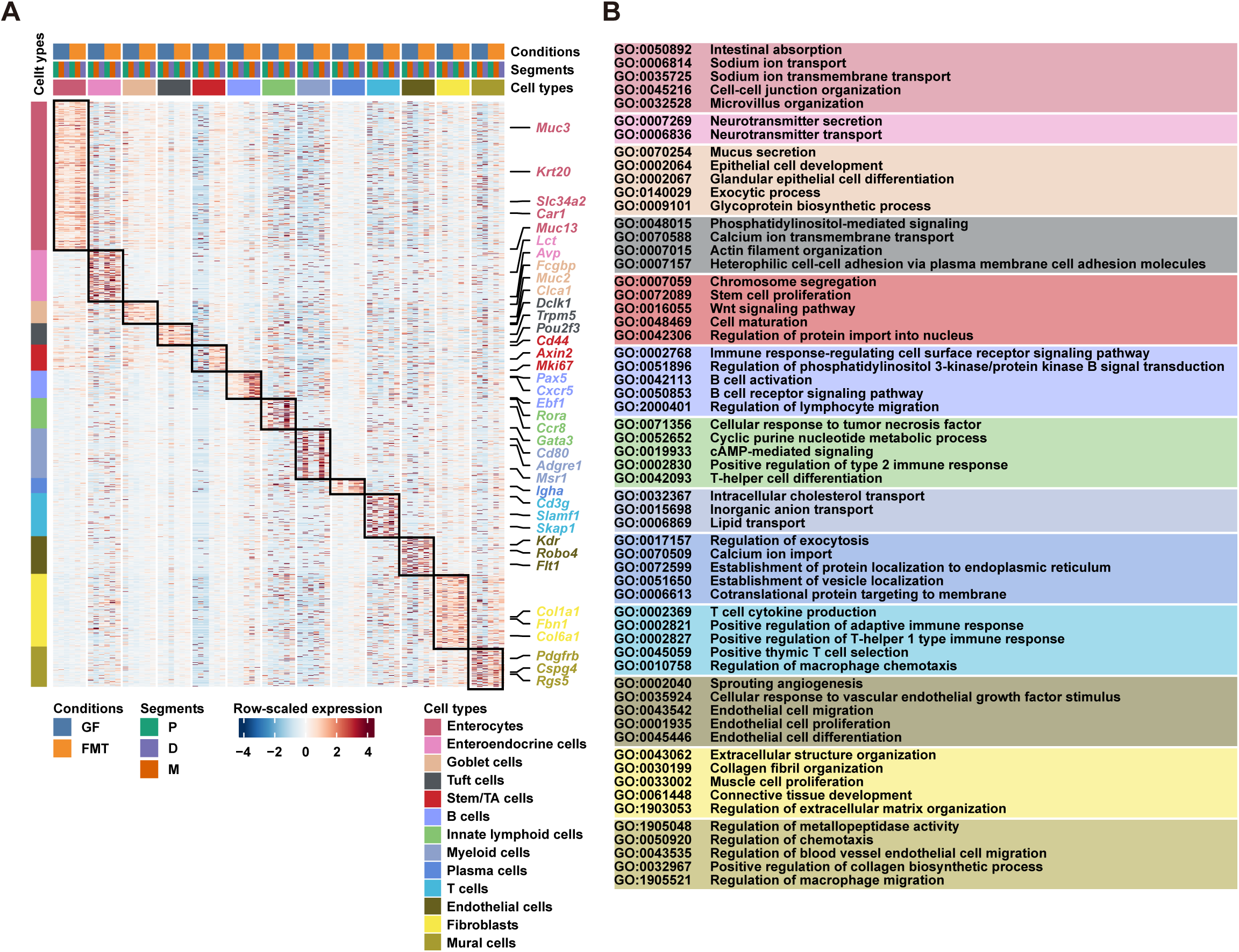
Transcriptional signatures and functional enrichment of distinct cell types. (A) Heatmap showing the row-scaled expression of up-regulated genes across distinct cell types and segments. Representative marker genes are indicated. (B) Representative GO biological pathways enriched for each annotated cell type.

**Extended Data Fig. 3.**
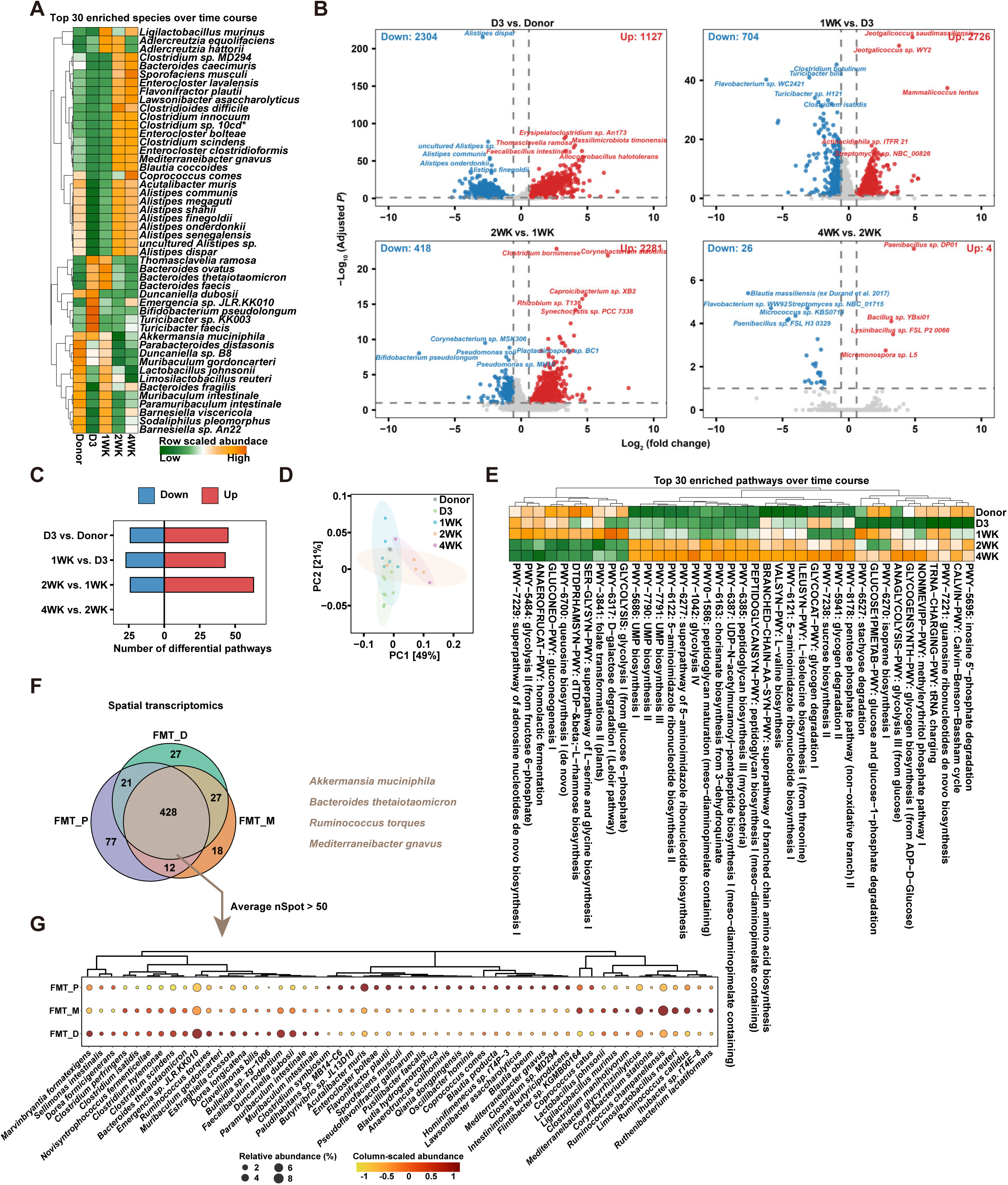
Temporal dynamics and regional distribution of microbial species. **(A)** Heatmap showing the temporal abundance changes of the top 30 enriched microbial species over the colonization time course. **(B)** Volcano plots showing the number of differentially abundant species across time point comparisons and compared to the donor. **(C)** Bar chart showing the number of differentially abundant metabolic pathways across time point comparisons. **(D)** Principal coordinate analysis (PCoA) based on pathway abundance illustrating the metabolic successional trajectory. **(E)** Heatmap showing the row-scaled abundance of the top 30 enriched metabolic pathways over the time course. **(F)** Venn plot showing the intersection of shared microbial species among the FMT segments. **(G)** Dot plot showing the column-scaled relative abundance of the shared differential species across colon segments. Only species with nSpots > 50 are shown.

**Extended Data Fig. 4.**
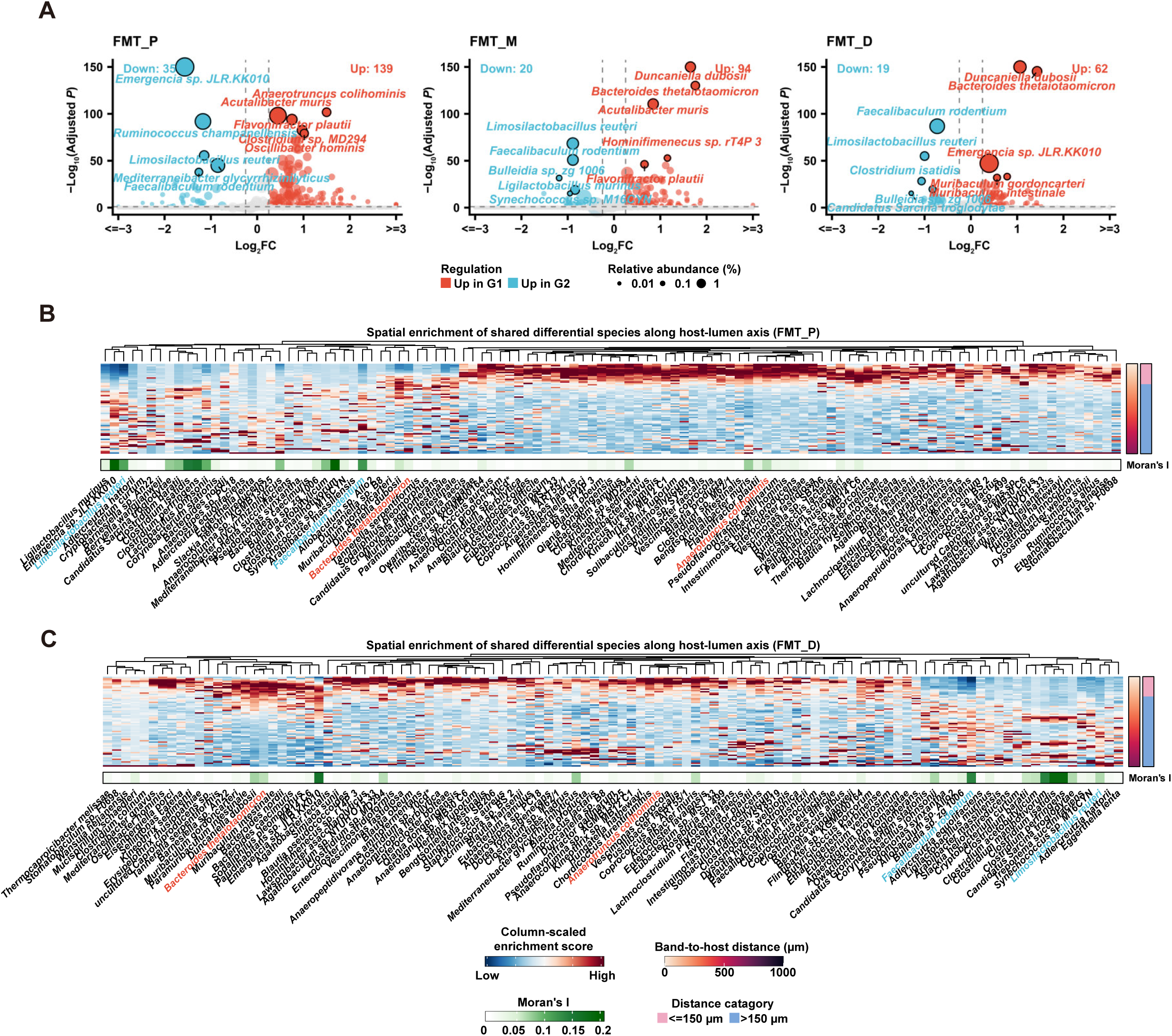
Spatial stratification of colonic microbes along the host– lumen axis. **(A)** Volcano plots highlighting differentially enriched microbial species between G1 (mucus-associated) and G2 (lumen-associated) regions. **(B-C)** Heatmaps showing the column-scaled enrichment scores of shared differential species along the host-lumen axis in FMT proximal (B) and distal (C) colons. Moran’s I, spatial autocorrelation index.

**Extended Data Fig. 5.**
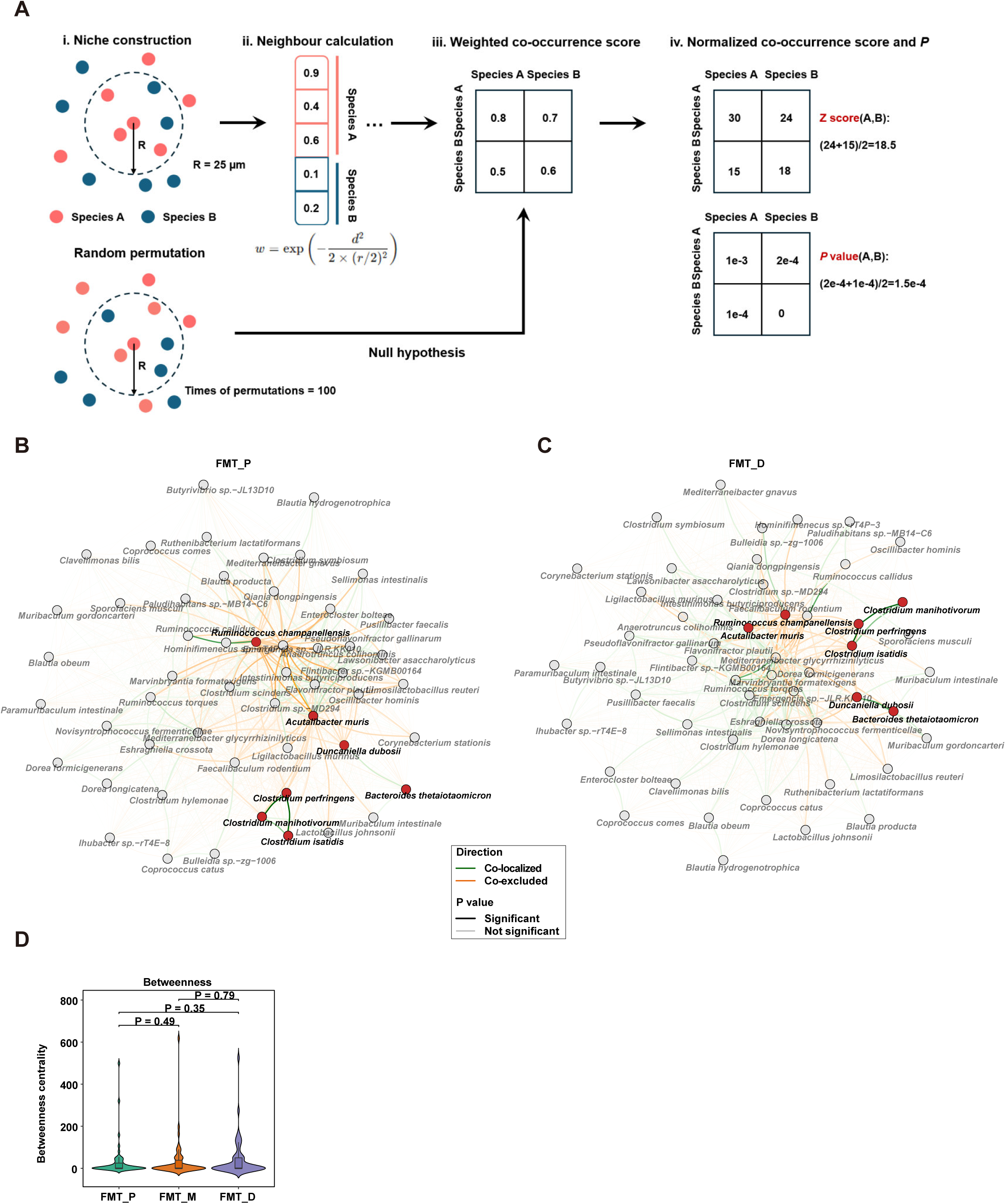
Spatial co-occurrence networks of microbial communities across colonic regions. **(A)** Schematic illustrating the computational framework for identifying spatial co-occurrence events, involving niche construction (i), neighbour calculation within a 25-µm radius (ii), calculating a weighted co-occurrence score (iii), and computing a normalized Z-score and P value based on random permutations (iv). **(B-C)** Spatial co-occurrence networks of microbial species in the FMT proximal (B) and distal (C) colon. Nodes represent species and edges indicate significant spatial co- localization (green) or co-exclusion (orange). Edge line type denotes statistical significance (*P* < 0.01). **(D)** Boxplots comparing the betweenness centrality of microbial species across colon segments. *P* values were calculated by a two-sided Kolmogorov–Smirnov test.

**Extended Data Fig. 6.**
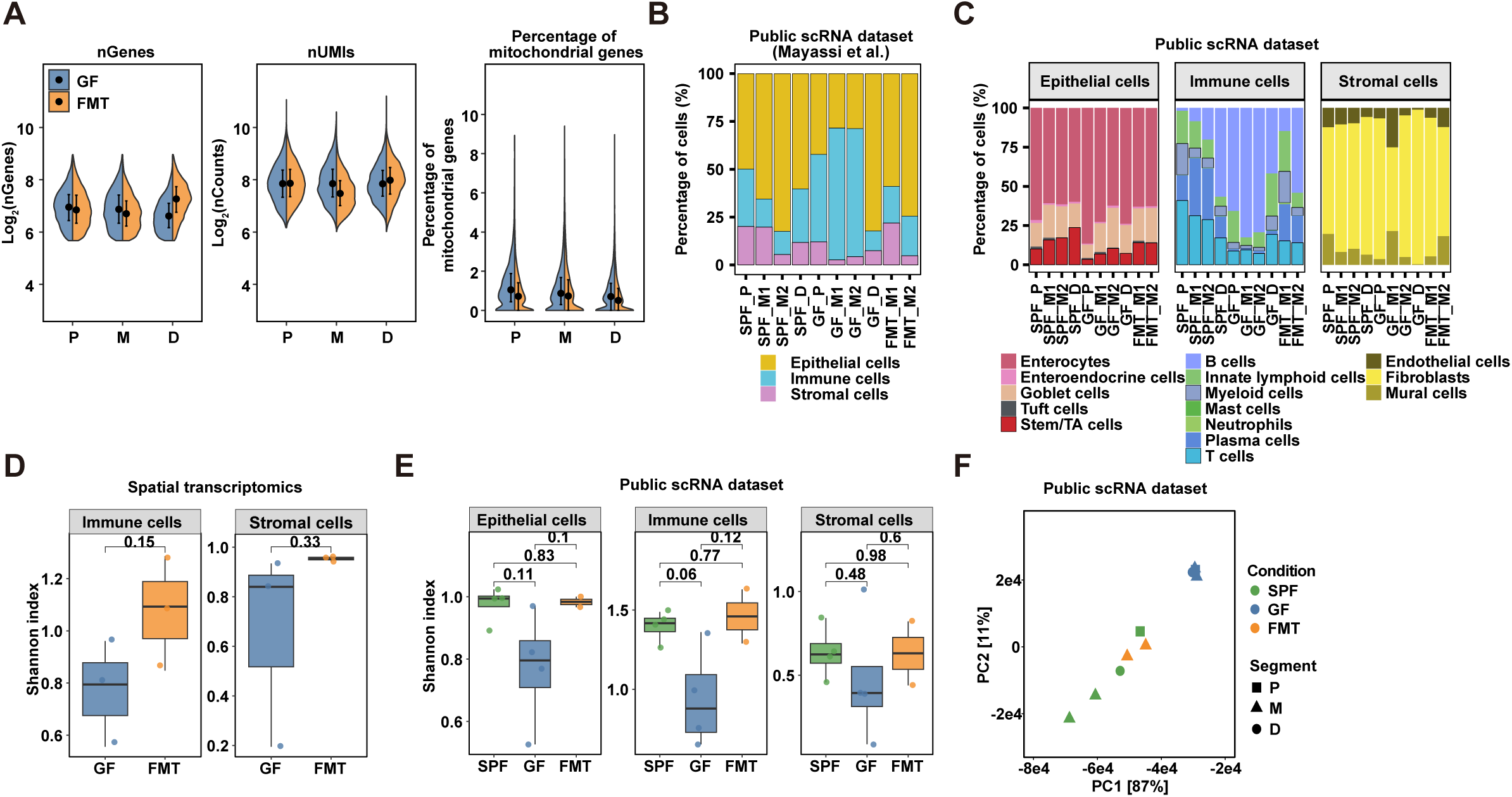
Quality control and cross-dataset validation of cellular composition and diversity analyses. **(A)** Violin plots showing quality control metrics of host spatial transcriptomics data, including the log_2_-transformed number of detected genes (left), log_2_-transformed UMI counts (middle), and percentage of mitochondrial genes (right) across colonic regions in GF and FMT mice. **(B)** Boxplots comparing the Shannon diversity index of immune (left) and stromal (right) cell populations in spatial transcriptomics data between GF and FMT mice. Student’s t- test. **(C-D)** Stacked bar charts showing the cellular composition of lineages (C) and cell types **(D)** in a o public scRNA-seq dataset (Mayassi et al.^13^). **(E)** Boxplots comparing the Shannon diversity index of epithelial, immune, and stromal lineages. **(F)** Principal component analysis (PCA) of pseudo-bulk gene expression of the published scRNA-seq dataset.

**Extended Data Fig. 7.**
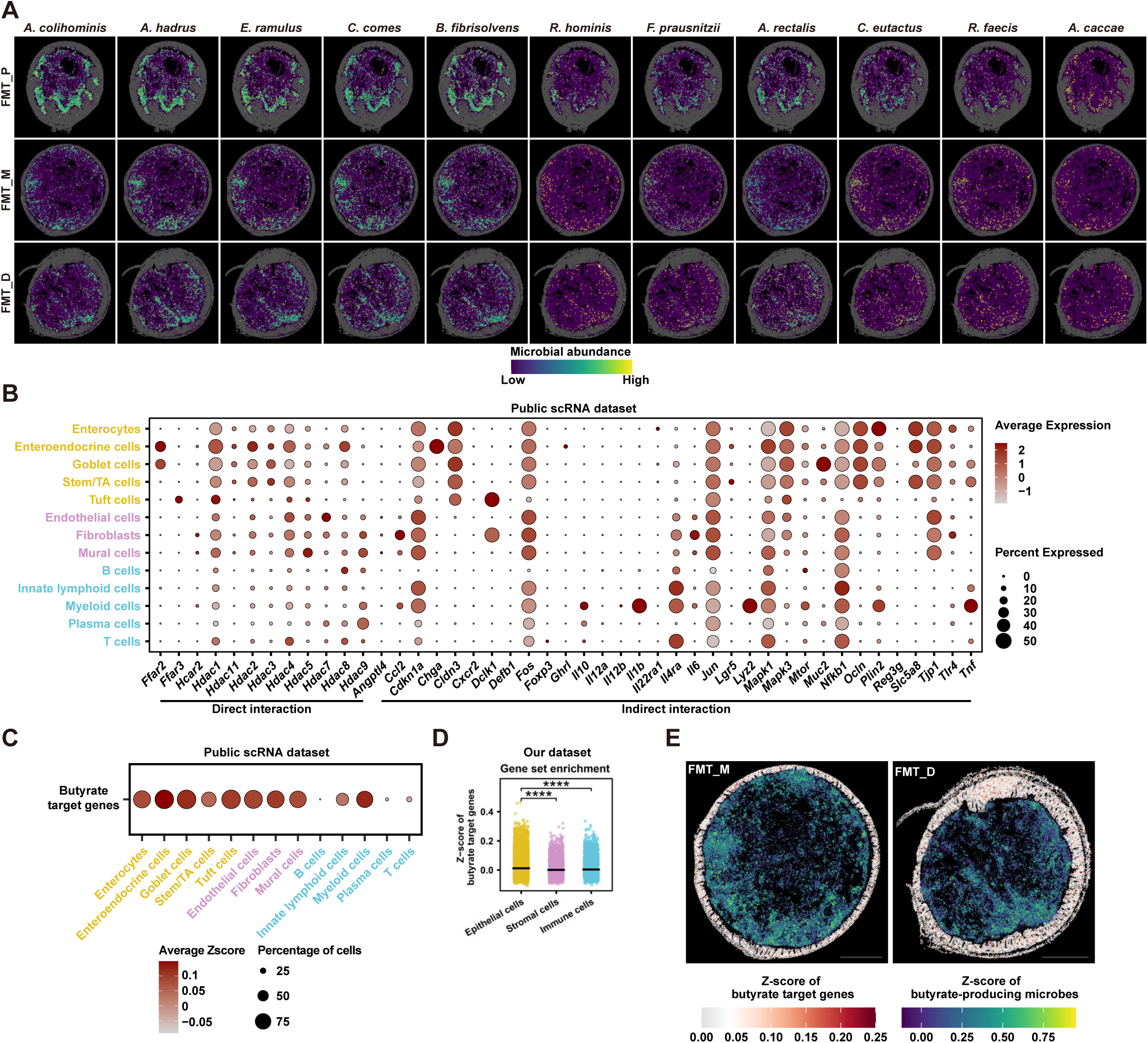
Spatial crosstalk of butyrate-producing microbes and host target gene expression. **(A)** Spatial maps showing the microbial abundance of up-regulated butyrate-producing species indicated in Fig. 6C. **(B)** Dot plot showing the average expression and percentage of cells expressing butyrate target genes (direct and indirect) across distinct cell types. **(C)** Dot plot showing the average Z-score of butyrate target genes and the percentage of expressing cells across distinct cell types. **(D)** Dot-line plot comparing the integrated butyrate target signature Z-score across epithelial, stromal, and immune cells. **** *P* < 0.0001; Wilcoxon rank-sum test. **(E)** Spatial maps of butyrate target gene expression and the abundance of butyrate-producing microbes from the FMT middle and distal segments.

